# Neuroprotective effects of Canagliflozin: lessons from aged genetically diverse UM-HET3 mice

**DOI:** 10.1101/2022.01.27.478017

**Authors:** Hashan S. M. Jayarathne, Lucas K. Debarba, Jacob J. Jaboro, Richard A. Miller, Marianna Sadagurski

**Author notes:** HSMJ and LKD contributed equally to this study. Corresponding author: Marianna Sadagurski, Department of Biological Sciences, Integrative Biosciences Center, Wayne State University, 6135 Woodward, Detroit, MI 48202.

## Abstract

The aging brain is characterized by a progressive increase in neuroinflammation and central insulin resistance, which contribute to neurodegenerative diseases and cognitive impairment. Recently, the Interventions Testing Program (ITP) demonstrated that the anti-diabetes drug, Canagliflozin (Cana), a sodium-glucose transporter 2 inhibitor (SGLT2i), led to lower fasting glucose and improved glucose tolerance in both sexes, but extended median lifespan by 14% only in male mice. Here we show that Cana treatment significantly improved central insulin sensitivity in the hypothalamus and the hippocampus of 30-month-old male mice. Remarkably, Cana-treated mice showed significant reductions in age-associated hypothalamic gliosis with a decrease in inflammatory cytokine production by microglia in both sexes. In the hippocampus, Cana reduced microgliosis and astrogliosis in males, but not in female mice. The decrease in microgliosis was partially dependent on mTOR signaling, as evidenced by reduced phosphorylation of S6 kinase in microglia of Cana-treated aged male, but not female mice. Remarkably, Cana treatment improved exploratory and locomotor activity of 30-month-old male but not female mice. Taken together, our findings demonstrate sex-specific neuroprotective effects of Cana treatment, suggesting its application for the potential treatment of neurodegenerative diseases.

## Introduction

The NIA-sponsored Interventions Testing Program (ITP) was designed to identify therapeutic interventions that slow aging in mice, as assessed by increased lifespan, and has identified several treatments that successfully extend lifespan in the genetically diverse UM-HET3 mouse model (Nadon *et al*. 2008; Harrison *et al*. 2014). The ITP recently reported that an FDA-approved anti-diabetes drug, Canagliflozin (Cana), a sodium-glucose transporter 2 inhibitor (SGLT2i), extended the median survival of male mice by 14%, without an effect on lifespan in females (Miller *et al*. 2020). UM-HET3 mice are not prone to diabetes, and most deaths are attributable to various forms of neoplasia, suggesting that tumorigenesis was inhibited or decelerated by this drug, though only in males. Interestingly, Cana led to lower fasting glucose and improved glucose tolerance in both males and females, and diminished fat mass in females only (Miller *et al*. 2020). Sex-specific lifespan benefit was also seen in male mice given another anti-diabetes drug, Acarbose, which also reduces blood glucose by slowing the breakdown of carbohydrates in the intestine (Lam *et al*. 1998; Harrison *et al*. 2014). However, it is yet to be determined whether reductions in blood glucose per se are responsible for lifespan extension.

Improving glucose regulation and metabolic control by pharmacological interventions has been shown to be beneficial in delaying some aspects of aging (Gonzalez-Freire *et al*. 2020). The cellular or molecular mechanism primarily responsible for this effect is not fully understood, although studies indicate that the interplay between lifespan extension and metabolic changes involves the ability to reduce systemic and/or central inflammation (Furman *et al*. 2019). For example, treatment of male mice with Acarbose or with 17-α-estradiol (17αE2), an optical isomer of 17-β-estradiol (Zhurova et al. 2009) that extends lifespan preferentially in male mice only (Strong *et al*. 2016), reduces age-associated metabolic and inflammatory dysfunction in the periphery and the brain (Garratt *et al*. 2017; Sadagurski *et al*. 2017). Similarly, the anti-diabetes drug Metformin exerts its neuroprotective effects in aging by reducing neuroinflammation (Kodali *et al*. 2021), although metformin appears not to increase lifespan in male or female mice tested by the ITP (Strong *et al*. 2016). Age-associated neuroinflammation triggered by activated microglia and astrocytes induces neuronal stress that affects brain insulin signaling, cognitive impairment, and progression of neurodegenerative diseases (von Bernhardi *et al*. 2015).

Limited studies link SGLT2i treatment with neuroprotection. For example, the SGLT2i Empagliflozin reportedly reduced beta-amyloid levels and improved cognitive abilities in a murine model of Alzheimer’s disease (AD) crossed to the diabetes model of leptin receptor deficiency (db/db), or in db/db mice alone (Hierro-Bujalance *et al*. 2020),(Lin *et al*. 2014). A recent clinical study demonstrated improved insulin sensitivity in the hypothalamus in subjects with prediabetes after treatment with Empagliflozin (Kullmann *et al*. 2021). Likewise, Cana reduced obesity-associated neuroinflammation in the hypothalamus and nodose ganglion (Naznin et al. 2017). However, the underlying mechanisms of the beneficial effect of the SGLT2i were associated primarily with diabetes and obesity. The question of whether Cana has more general beneficial effects on the aging brain or protective effects from age-associated neurodegeneration, in diabetes-free mice, remains open. In this study, we evaluated the possible neuroprotective effect of long-term Cana treatment in aged UM-HET3 mice, with a specific focus on brain regions that are sensitive to metabolism and cognitive function.

## Results

### Canagliflozin treatment improves central insulin sensitivity in aged mice

We have previously demonstrated that UM-HET3 male and female mice treated =with Cana from 7 months of age had a significantly lower fasting glucose level and showed an improved glucose tolerance by 22 months of age (Miller *et al*. 2020). To investigate whether Cana treatment affected central insulin sensitivity, we administered insulin intraperitoneally and assessed the distribution of FoxO1 transcription factor between the nuclear and the cytoplasmic compartments, in the hypothalamus and the hippocampus of 30-months-old control and Cana-treated mice after 23-months of Cana treatment. We found that long-term Cana treatment sensitized the hypothalamic response to insulin, as shown by increased levels of cytoplasmic FoxO1 in the arcuate nucleus (ARC) of the hypothalamus in response to insulin administration in both sexes. In contrast, old control males exhibited insulin resistance (sex x insulin interaction p<0.01) (Figure 1A, B G and H). A similar pattern in response to insulin stimulation was observed in the CA3 and DG sub-regions of the hippocampus, with a significant interaction effect between Cana and insulin only in males (sex x insulin interaction p<0.0001) (Figure 1C, E, and G). Both control and Cana-treated females responded similarly to insulin stimulation with elevated cytoplasmic levels of FoxO1 in the hypothalamus and hippocampus without significant effect of Cana (Figure 1B, D, F, and H). These results are in accordance with previous reports indicating sex-specific sensitivity to insulin action in the brain on food intake and body weight in humans and rodents (Clegg *et al*. 2003; Hallschmid *et al*. 2004), demonstrating protective properties of Cana treatment in the brain beyond its effects on peripheral metabolism.

**Figure 1:**
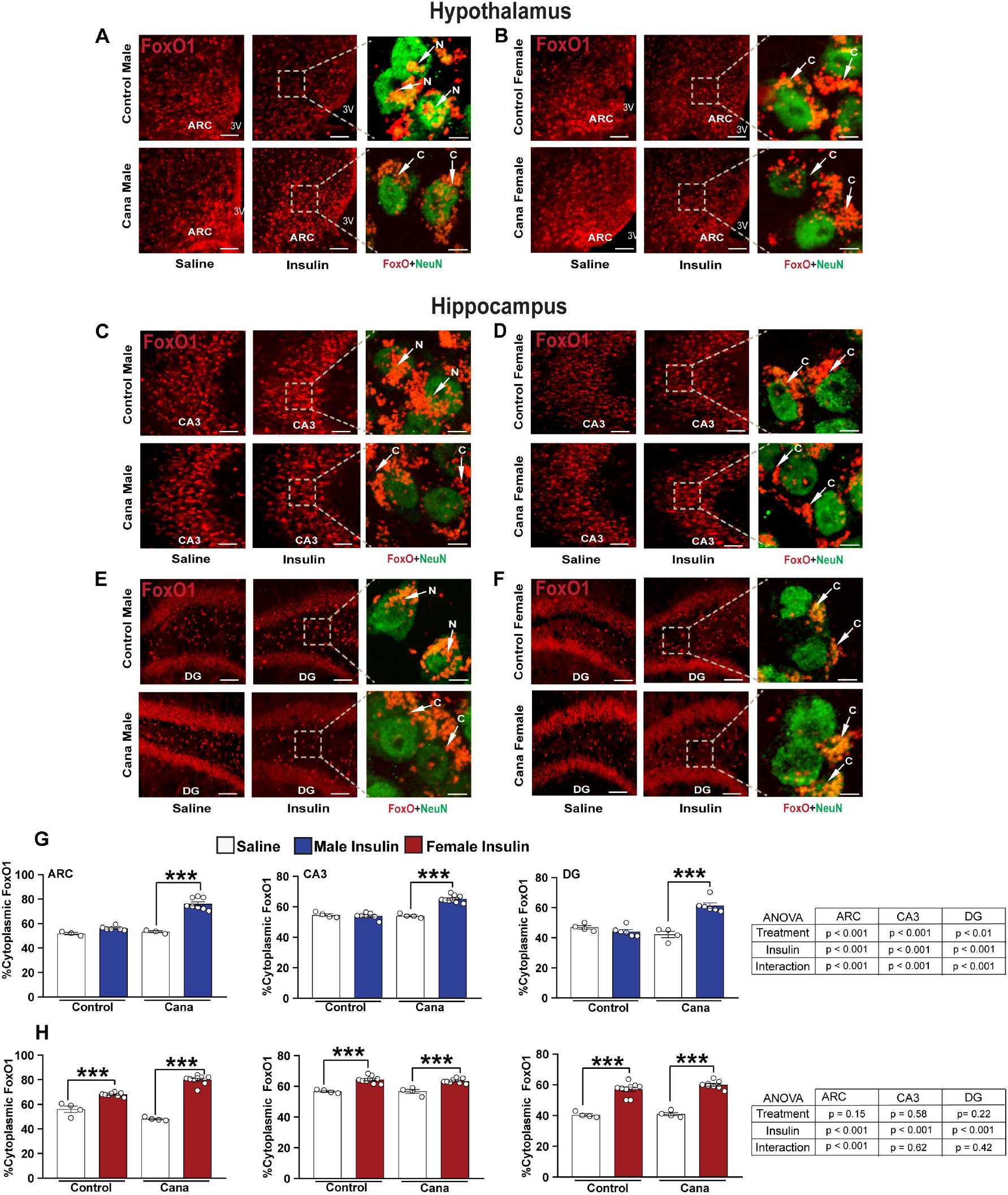
Insulin→FoxO1 signaling in aged Cana treated mice. Immunofluorescence for FoxO1 (red) and NeuN (green) in 30-months-old control and Cana treated mice injected with insulin (3 U/kg i.p.; 15min) or saline. Representative images from the hypothalamus in males (A) and females (B), or hippocampus CA3 in males (C) and females (D), and dentate gyrus (DG) in males (E) and females (F) of control and Cana-treated mice are shown. Scale bars represent 100 μm or 5 μm for the left panels for confocal images of cells from the boxed areas from the right panel, showing FoxO1 (red) merged with NeuN (green) (right) in 30-months-old mice of the indicated groups. White arrows indicate the localization of the FoxO1, C. cytoplasmatic, N. nuclear. 3V, third ventricle. Quantification of neurons containing cytoplasmic FoxO1 immunoreactivity in the ARC, CA3, DG in males (G) and females (H); error bars show SEM for n = 4-7 mice/group. Images of matched brain areas from each mouse were taken from at least “three-four” sections containing the hypothalamus and hippocampus. Data were analyzed by 2-factor ANOVA and further analyzed with the Newman-Keuls post hoc test (***p<0.001). The tables demonstrate the two-factor ANOVA analysis.

### Cana reduces the mTOR signaling in a sex-specific manner in the hippocampus

Brain insulin resistance is associated with hyperactivation of the mechanistic target of rapamycin (mTOR), but inhibition of the mTOR complex 1 (mTORC1) increases animals’ lifespan (Lamming *et al*. 2012). We assessed the phosphorylation state of S6 kinase (pS6 S240/S244), a downstream substrate of mTORC1, in the hypothalamus of 30-month-old mice treated with Cana. We found a significant effect of sex, with females expressing higher levels of phosphorylated S6 (pS6) than males (p<0.001), but there was no effect of Cana treatment on mTORC1 (Figure 2A and B). We detected significant sex-specific effects in mice fed the control diet, where females expressed higher levels of pS6 in the hypothalamus (p<0.01), while males expressed higher pS6 in the hippocampal CA3 sub-region as compared to females (p<0.001). In the CA3 and DG sub-regions the levels of pS6 were significantly reduced in Cana-treated male but not in female mice (Figure 2C and D), with a significant interaction effect between treatment and sex (p<0.001 for CA3 and p<0.05 for DG). Thus in the hippocampus, sex differences on pS6 were eliminated by Cana treatment.

**Figure 2:**
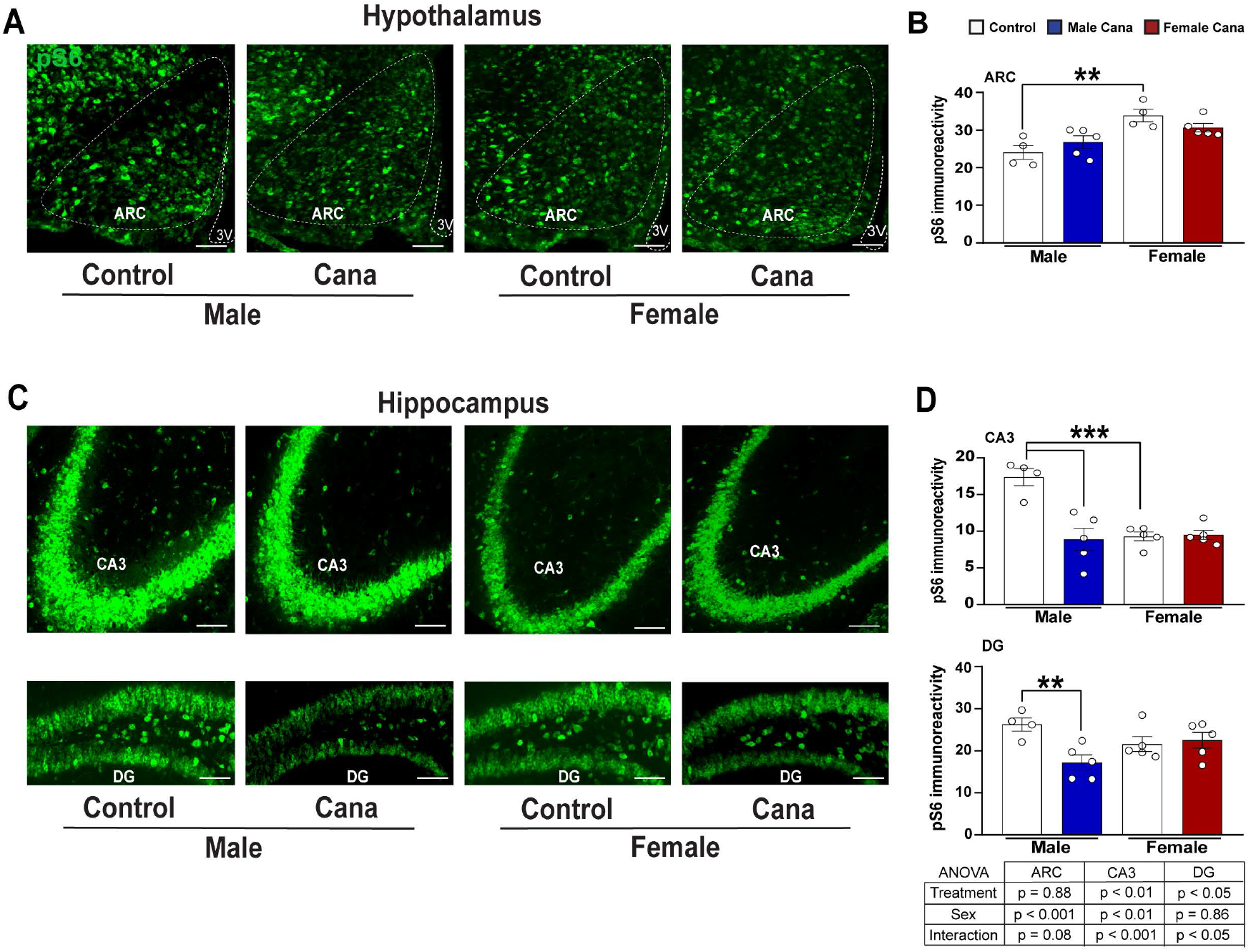
pS6 protein expression in aged Cana treated mice. Brain sections of 30-months-old male and female mice were analyzed for hypothalamic and hippocampal phosphorylated S6 protein expression. Representative images showing immunostaining in the arcuate nucleus of the hypothalamus (ARC) (A), and hippocampal CA3, and dentate gyrus (DG) (C) of control and Cana-treated male and female mice. Quantification of pS6 immunoreactivity in the ARC (B), CA3 and DG (D) of control and Cana-treated male and female mice. Scale bars: 200 μm, 3V, third ventricle; error bars show SEM for n = 4-5 mice/group. Images of matched brain areas from each mouse were taken from at least “three-four” sections containing the hypothalamus and hippocampus. Data were analyzed by 2-factor ANOVA and further analyzed with the Newman-Keuls post hoc test (**p<0.01, ***p< 0.001). The tables demonstrate the two-factor ANOVA analysis.

### Cana treatment reduces neuroinflammation in a region- and the sex-specific manner in aged mice

Activation of microglia and astrocytes is fundamental for neuroinflammation, which is in turn tightly linked with insulin resistance in the aging brain (Komleva *et al*. 2020). We assessed whether Cana treatment modulated inflammatory cytokine production by microglia and astrocytes in the aging brain. Microglia and astrocytes were purified from 30-month old mice by density gradient followed by cell sorting of whole brains (Figure 3A and B). Cells obtained by density gradient separation were then stained with CD45/CD11b and sorted to obtain populations enriched for CD11b^+^ /CD45^low^ cells, markers characteristics of resident brain microglia, and CD11b^+^/CD45^hi^ cells, which represent systemic macrophage populations. Astrocytes were sorted by ACSA2, an astrocyte surface antigen (Kantzer *et al*. 2017). Microglia and astrocyte preparations were verified by expression of microglia-specific or astrocyte-specific genes, respectively (Supplemental Figure 1A, B). Cana treatment significantly reduced the expression of mRNA for the inflammatory cytokines *TNFα, IL6*, and *IL1* in microglia in both male and female mice as compared to control microglia, with an interaction effect between Cana and sex only for *IL6* gene expression (p<0.01) (Figure 3A). Further, Cana treatment significantly reduced the expression of the neuroinflammatory A1-reactive genes, *Ligp* and *H2t23*, in astrocytes (Figure 3B). *IL33 is* a cytokine that mediates hippocampal synaptic plasticity and was shown to increase in astrocytes with age (Carlock *et al*. 2017; Wang *et al*. 2021). Interestingly, the expression of *IL33* by astrocytes was reduced by Cana treatment only in males but not in females (sex x treatment interaction p<0.001) (Figure 3B).

**Figure 3:**
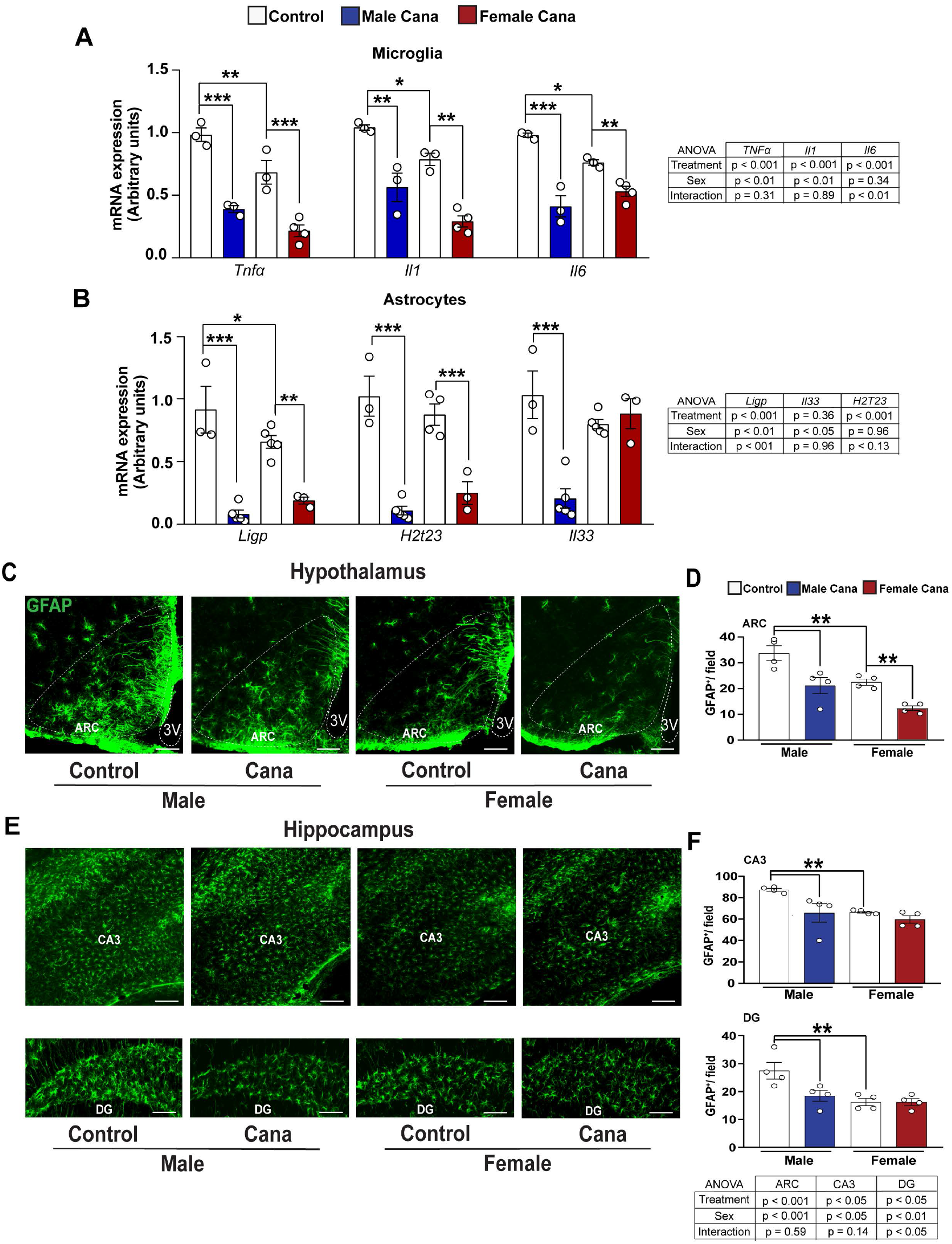
Neuroinflammation in aged Cana treated mice. Expression of *Tnfa, Il1*, and *Il6*, from isolated microglia (A) and *Il33, Ligp* and *H2t23* from isolated astrocytes (B) of 30-months-old control and Cana-treated male and female mice. Error bars show SEM for n = 3-5 mice/group. Data were analyzed by 2-factor ANOVA and further analyzed with the Newman-Keuls post hoc test (*p < 0.05, **p<0.01, ***p< 0.001). Brain sections of male and female mice were analyzed for hypothalamic and hippocampus GFAP^+^ astrocytes. Representative images showing immunostaining in the arcuate nucleus of the hypothalamus (ARC) (C), CA3, and dentate gyrus (DG) hippocampus areas (E) of 26-28 months-old control and Cana-treated mice. Scale bars: 200 μm. 3V, third ventricle. Numbers of cells immunoreactive for GFAP in the ARC (D), CA3 and DG (F) from indicated male and female mice; error bars show SEM for n = 4 mice/group. Images of matched brain areas from each mouse were taken from at least “three-four” sections containing the hypothalamus and hippocampus. Data were analyzed by 2-factor ANOVA and further analyzed with the Newman-Keuls post hoc test (**p<0.01). The tables demonstrate the two-factor ANOVA analysis.

To assess the region- and sex-specific effects of Cana on neuroinflammation we examined the activation state of astrocytes and microglia in aged brains. We evaluated the effect of Cana on astrogliosis in both the hypothalamus and hippocampus using glial fibrillary acidic protein (GFAP) as a marker for the astrocytes. In aged mice, we found about 1.5-fold lower numbers of hypothalamic astrocytes in females (p<0.001) as in our previous studies using younger mice, 12 or 22 months of age (Sadagurski *et al*. 2017). We similarly detected lower numbers of the astrocytes in the hippocampus of female mice as compared to males (p<0.05). Cana treatment reduced astrogliosis in the hypothalamic ARC in both male and female mice (Figure 3C and D). In the hippocampus, Cana reduced astrogliosis in the sub-regions CA3 and DG in males, but not in female mice (Figure 3E and F), with a significant interaction effect between drug (treatment) and sex in DG (p<0.05). In addition, both male and female mice treated with Cana showed a significantly reduced number of hypothalamic microglia cells positive for the ionized calcium-binding adapter molecule 1 (Iba1), a microglia-specific marker (Figure 4A and B). The same effect was also observed by staining with TMEM119, another microglia marker (Supplementary Figure 2). Hypothalamic Iba1^+^ cells produced tumor necrosis factor-alpha (TNF-α), indicating an inflammatory state (Figure 4C and D). Cana significantly reduced TNFα, production by microglia in both sexes in the ARC (p<0.001) (Figure 4D). Similar to the hypothalamic astrogliosis, we found a significant effect of sex on hypothalamic microgliosis (p<0.05), with lower microglia numbers in females compared to male mice. Sex effect was also observed on microglia numbers in the hippocampus (Figure 4E and F). Cana reduced microgliosis in sub-regions CA3 and DG in males, but not in female mice (Figure 4E and F), with a significant interaction effect between drug and sex only in CA3 (Figure 4F, p<0.01). Cana had no effect on microglial numbers in the female hippocampus in both CA3 and DG regions (Figure 4E and F). We were not able to reliably detect TNFα by immunostaining in the hippocampus (data not shown). Overall, these data indicate that Cana supplementation reduces age-associated neuroinflammation in regions critical for the integration of metabolic homeostasis and memory formation.

**Figure 4:**
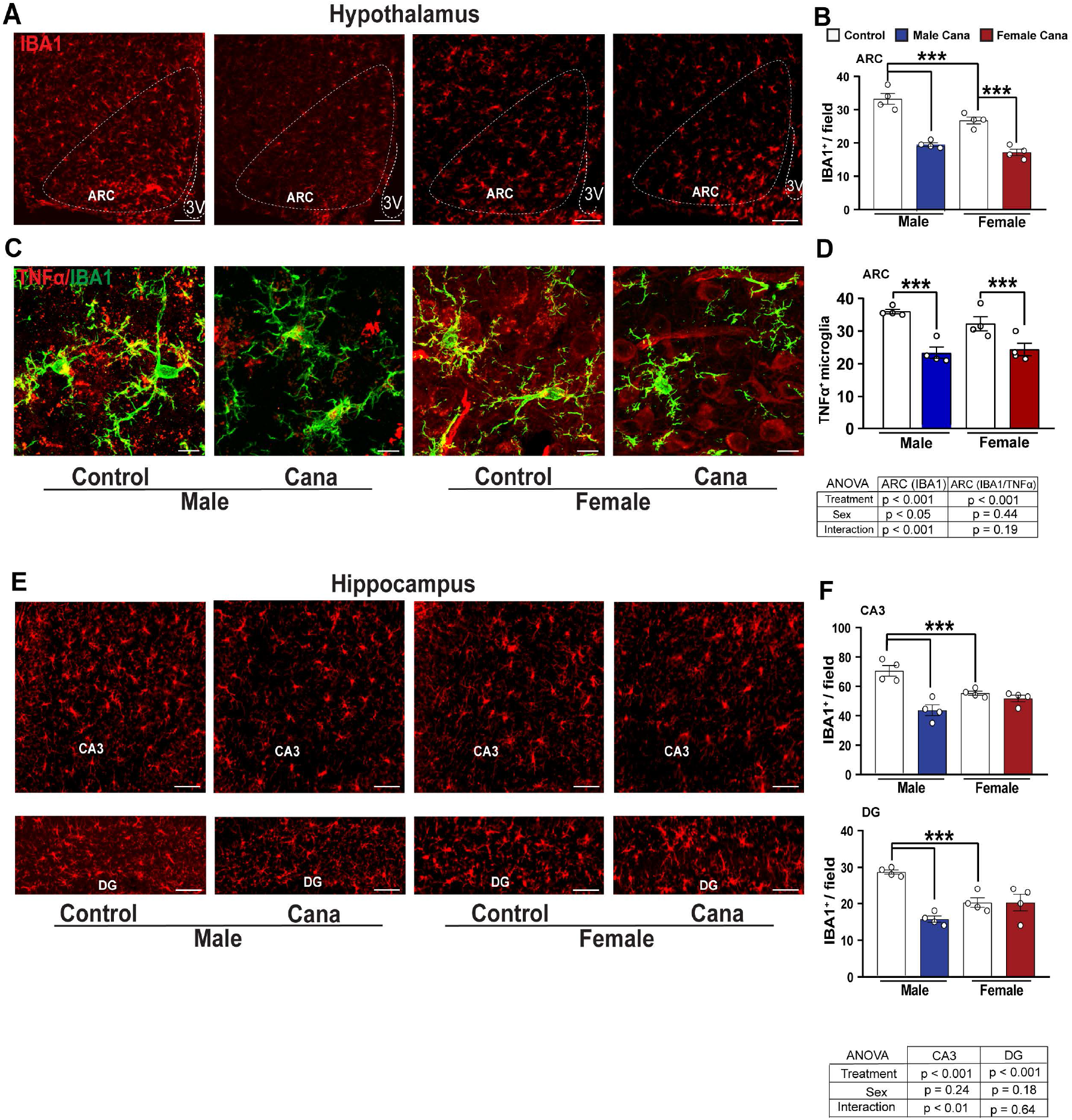
Microgliosis in aged Cana treated mice. Representative images showing Iba1^+^ cells (red) (A) and confocal images showing Iba1 (green) merged with TNF-α (red) (C) immunostaining in the arcuate nucleus of the hypothalamus (ARC) of control and Cana-treated 26-28 months-old male and female mice. Numbers of cells immunoreactive for (B) Iba1 and (D) TNF-α^+^/Iba1^+^ in the ARC of control and Cana-treated male and female mice. (E) Representative images showing Iba1^+^ cells (red) in the CA3, and DG of 26-28 months-old control and Cana treated mice. Numbers of cells immunoreactive for Iba1 in the CA3 and DG (F) from indicated male and female mice; error bars show SEM for n = 4 mice/group. Images of matched brain areas from each mouse were taken from at least “three-four” sections containing the hypothalamus and hippocampus. Scale bars: 200 μm, 10 μm on merged images. 3V, third ventricle. Data were analyzed by 2-factor ANOVA and further analyzed with the Newman-Keuls post hoc test (***p < 0.001). The tables demonstrate the two-factor ANOVA analysis.

### Cana treatment reduces mTOR signaling in aged microglia in a sex-specific manner

Microglia from aged mice demonstrated upregulation of mTOR-downstream signaling and the production of inflammatory cytokines (Keane et al. 2021). Interestingly, we detected sexspecific differences in mTOR signaling, as indicated by phospho-S6 (pS6) in aged microglia, with females exhibiting lower phospho-S6 levels in both hypothalamus (p<0.05) and hippocampus (p<0.001 for CA3). pS6 is lowered in hypothalamic microglial cells in response to Cana treatment in both male and female mice with a stronger effect in male mice (Figure 5 and Supplementary Figure 3). The hippocampus, however, is different: the fraction of pS6 positive microglia cells was significantly reduced in the hippocampus in response to Cana only in males (for an interaction effect between treatment and sex p<0.0001 for CA3 and p<0.05 for DG) (Figure 5C and D). This sexually dimorphic effect of Cana on mTOR signaling in microglia is consistent with the male-specific lifespan extension observed in Cana-treated animals (Miller *et al*. 2020).

**Figure 5:**
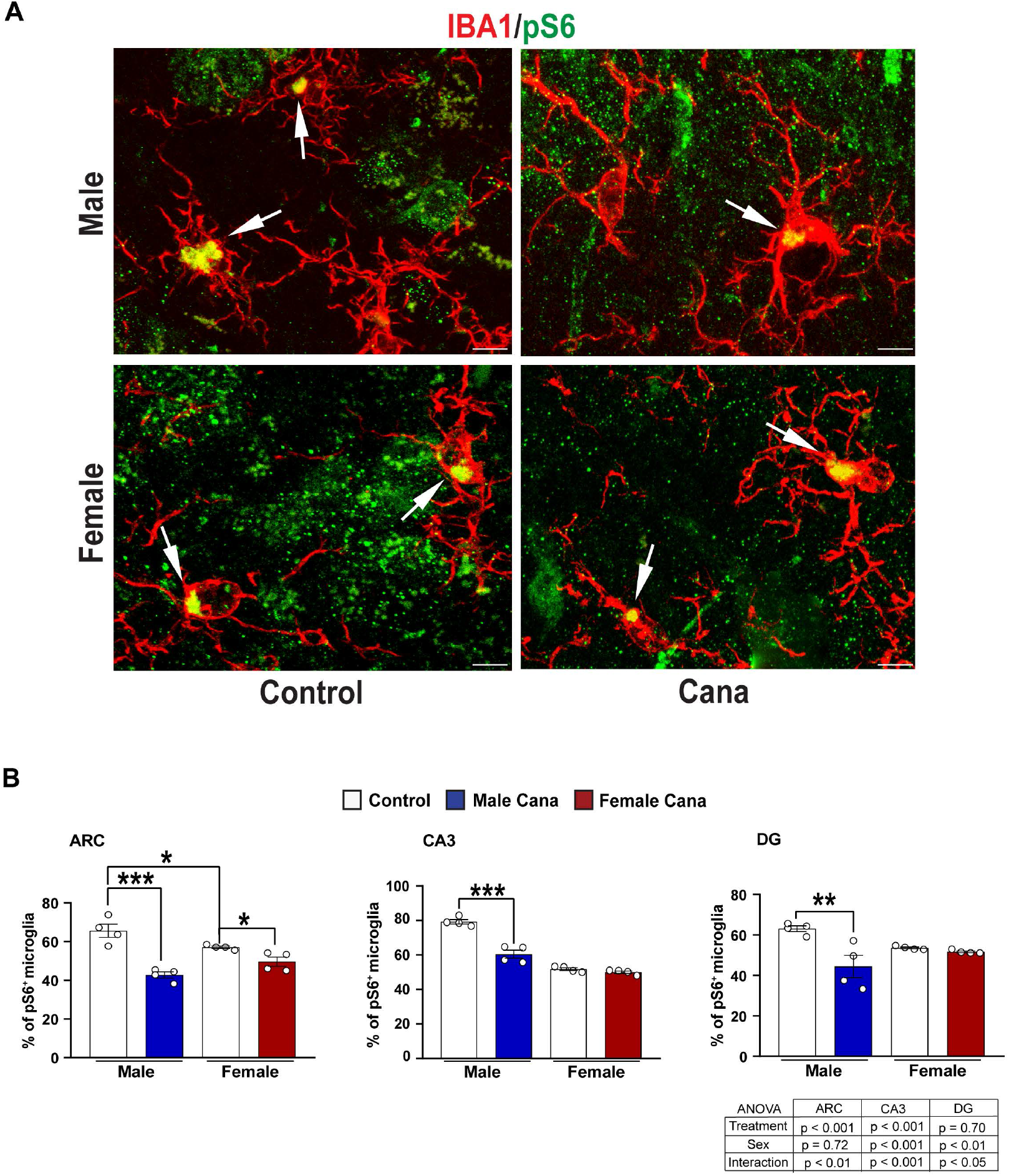
Microglial pS6 expression in Cana treated aged mice. (A) Representative confocal images of phosphorylated S6 (pS6, green) and Iba^+^ cells (red) in the arcuate nucleus of the hypothalamus (ARC) of 26-28 months-old control and Cana treated male and female mice. Scale bars: 5 μm, 3V, third ventricle. (B) Percentage of Iba1^+^ pS6 in the ARC, CA3, and DG of male and female mice. Error bars show SEM for n = 4 mice/group. Data were analyzed by 2-factor ANOVA and further analyzed with the Newman-Keuls post hoc test (**p < 0.01; ***p < 0.001). The tables demonstrate the two-factor ANOVA analysis. Images of lower magnification in hypothalamus and hippocampus are shown in Supplementary Figure 3.

### Cana treatment improves locomotor activity and overall behavior function in aging

To assess neuromuscular function, we performed Rotarod tests and grip strength analyses. We assessed rotarod performance at 22 months of age using an acceleration protocol where mice were tested for their ability to balance on a progressively accelerating rotarod (Garratt *et al*. 2019). The ability of mice to maintain balance on the rod declines with age, while Cana treatment significantly improves balance ability and was associated with a longer time on the accelerated rotarod (mean of 3 trials) before fall (p = 0.011) independent of sex (Figure 6A). Similarly, the Cana effect on the maximum performance on the rotarod was increased (p = 0.025), while no interaction with sex was detected (Figure 6B). Forepaw grip score did not differ significantly between Cana-treated or untreated mice at this age (Figure 6C). Body weight was significantly decreased by about 9 grams in Cana-treated females and about 7 grams in males (45.92±1.86 for male control versus 39.05±1.14 for male Cana; 43.56±1.92 for female control versus 34.65±1.08 for female Cana), indicating that improved rotarod performance may be due, at least in part, to lower body weight in the Cana groups. To evaluate general activity levels, gross locomotor activity, and exploratory behavior, we performed an open field test in 30-month-old control and Cana-treated mice. Cana-treated male mice were significantly more active than control males as shown by the increased total distance traveled (Figure 6D, E, and F). The latency to enter the center of the arena and the numbers of visits and duration in the center were significantly increased in males as well (Figure 6G). There was a significant interaction between drug and sex (p<0.05). A significantly increased duration in the center of the arena of Cana treated male mice, compared to control males, suggested reduced anxiety-related behavior, and increased willingness to explore a new environment (Shoji *et al*. 2016). None of these measures was altered by Cana in females (Figure 6D-H).

**Figure 6:**
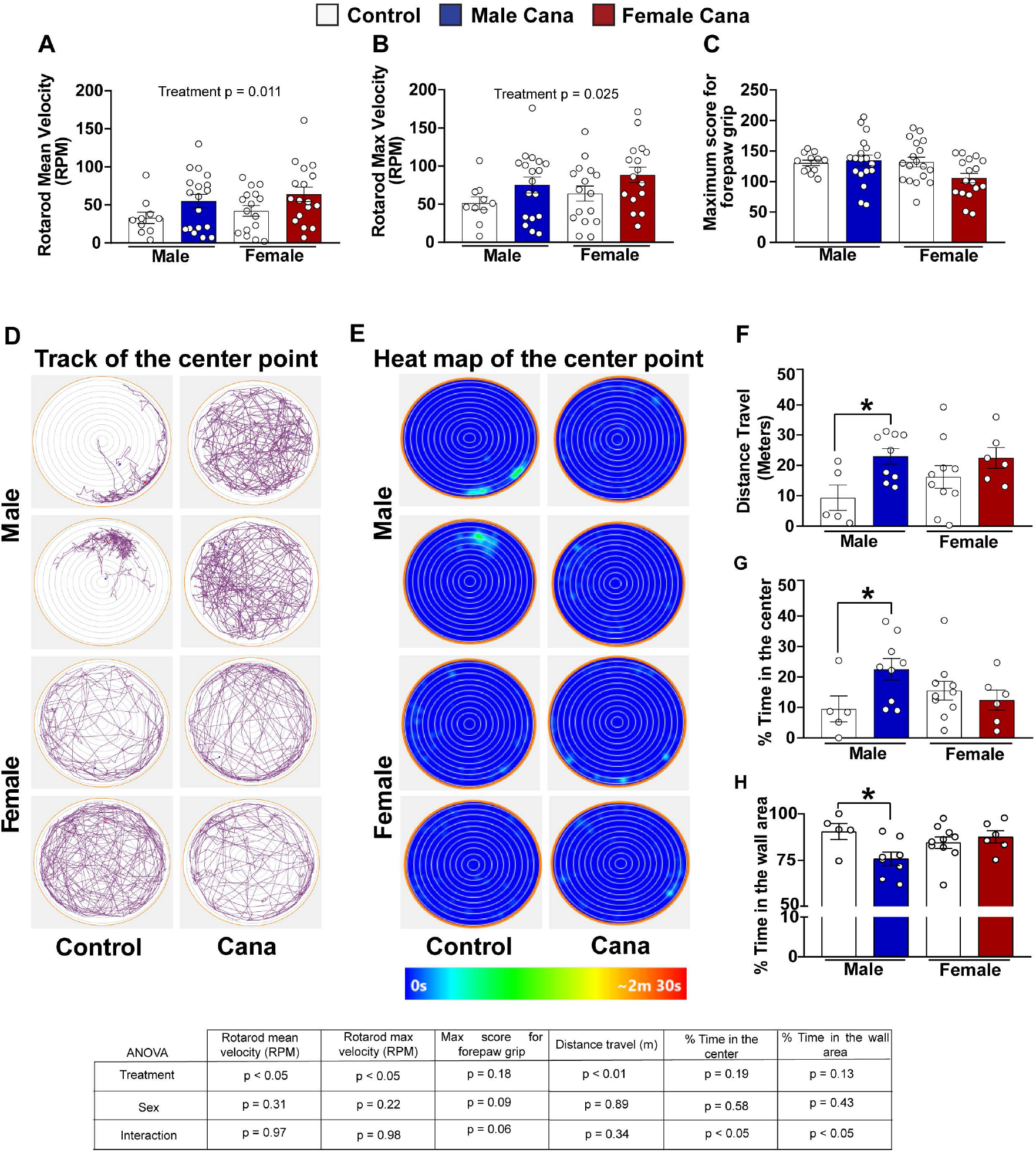
Neuromuscular function, locomotor activity, and exploratory behavior in Cana treated aged mice. (A) Rotarod mean velocity (RPM), (B) Maximum velocity, and (C) Maximum score for forepaw grip of 22-month-old control and Cana treated male and female mice. Error bars show SEM for n = 12-19 mice/group. Representative images from the open field test (OFM) of 30 months-old control and Cana treated male and female mice. (D) Track of the center point represent the total distance traveled by the mice during 10 min; (E) Heat map of the center point, the light-blue color represents the time spent during 10 min in the same spot. (F) Total distance traveled (meters) in the OFM; (G) Percentage time spent in the center of OFM; (H) Percentage time spent in the outer wall area. Error bars show SEM for n = 5-9 mice/group. Two-factor ANOVA analyzed data and further analyzed with the Newman-Keuls post hoc test (*p < 0.05). Tables show the two-factor ANOVA analysis.

## Discussion

We identified a broad spectrum of cellular and functional changes that were improved by Cana treatment in genetically heterogeneous aged mice. Cana treatment significantly improved central insulin sensitivity, reduced age-associated gliosis, and decreased microglial mTOR signaling in a sex-specific manner, predominantly benefiting old male but not female mice. Measures of physical performance were significantly improved by Cana in male but not in female mice. We posit that Cana, and potentially other SGLT2 inhibitors, protect against age-dependent neurological, behavioral, and metabolic decline even in the absence of diabetes.

Cana treatment improved central insulin sensitivity in the hypothalamus and hippocampus of aged UM-HET3 male mice, while females maintained adequate insulin signaling regardless of treatment. In our previous study, we showed that Cana levels in brain extracts and plasma were significantly higher in females than in male mice (Miller *et al*. 2020). Thus, Cana-induced brain insulin sensitivity in males cannot be attributed to the higher systemic or central levels of the drug. A recent clinical study with the SGLT2i empagliflozin reported improved hypothalamic insulin sensitivity in subjects with prediabetes and obesity (Kullmann *et al*. 2021), although the study was not randomized in a sex-stratified manner. The mechanisms for improved central insulin sensitivity are still not fully clear. However, these authors speculated that SGLT2i can modulate the activity of the autonomic nervous system with a shift from sympathetic to parasympathetic tone (Kullmann *et al*. 2021). The current data, showing Cana-induced improvement in central insulin signaling in male mice, implies beneficial effects of Cana beyond those related to systemic glucose control per se. Indeed, insulin resistance in the brain has been associated with acceleration of the aged-induced cognitive decline in animal models and humans (Freude *et al*. 2009; Talbot *et al*. 2012). Sex-specific effects in brain insulin sensitivity may underlie or partially contribute to the benefit of Cana on lifespan in males.

Neuroinflammation, driven by activation of glial cells, increases with age in mouse and human brains and is associated with decreased central insulin signaling (Sierra *et al*. 2007). Interaction between inflammation and insulin sensitivity is well documented for many tissues, and is beginning to be appreciated in the CNS as well (Milstein & Ferris 2021). We show that Cana treatment significantly reduced microglia and astrocytes activation in a sex-and region-specific manner. In aged male mice treated with Cana, we found reduced microgliosis and astrogliosis in both the hypothalamus and hippocampus, while in female mice we found reductions in neuroinflammation only in the hypothalamus. This was evidenced by significantly reduced expression of the pro-inflammatory cytokines in microglia with Cana treatment. Age-related increases in pro-inflammatory cytokines in the brain are thought to be detrimental to humans or rodents and correlate with deficits in cognitive function (Barrientos et al. 2015). Aged-induced microgliosis promotes activation of A1-like reactive astrocytes that in turn upregulate many mediators with neurotoxic function, causing neuronal death and neurodegeneration (Liddelow et al. 2017; Clarke et al. 2018). Ample evidence suggests that impaired peripheral metabolism associated with hypothalamic inflammation contributes to impairments in insulin sensitivity (Jais & Bruning 2017). Our previous study revealed improved peripheral glucose tolerance and lower body weight in aged male and female mice with Cana treatment (Miller *et al*. 2020). In the current study, we show that Cana reduced hypothalamic gliosis in both sexes, which may contribute to improved whole-body metabolism. Reduced hippocampal gliosis in Cana-treated male, but not female mice, is consistent with the sex-specific lifespan increase seen in males and might be among the principal mechanisms by which Cana can improve cognitive function in aging.

Among commonly used SGLT2i, Cana has the least specificity and can potentially inhibit both SGLT1 and SGLT2 transporters (Ohgaki *et al*. 2016). SGLT1 and SGLT2 have been detected in many areas of the brain (Głuchowska *et al*. 2021). Brain expression of SGLT2 is lower than SGLT1 but was detected in the microvessels of the blood-brain barrier (BBB), hippocampus pyramidal and granular cells, and astrocytes in the ventromedial hypothalamus (Poppe *et al*. 1997; Fan *et al*. 2015; Tahara *et al*. 2016; Nguyen *et al*. 2020). Thus, while Cana’s effects on CNS can be indirect, secondary to peripheral changes, or mediated via the activity of autonomic inputs to the hypothalamus (Spallone & Valensi 2021), it is plausible that Cana can attenuate hypothalamic gliosis directly via binding to the SGLT1 or SGLT2. Additional studies will be required to address this question.

Cana treatment was associated with reduced mTOR signaling in the hippocampus of 30 month-old male mice, as shown by reduced phosphorylation of ribosomal protein S6, one of the major downstream targets of the mTOR pathway. Previous studies have shown that Cana’s effects on metabolic health were associated with suppressed mTOR signaling, as shown by reduced hepatic pS6 signaling accompanied by increasing AMPK activity in young animals (Osataphan et al. 2019). That study did not, however, evaluate the effects of Cana treatment in female mice. The age trajectory of mTORC1 (mTOR complex 1) and mTORC2 (mTOR complex 2) activity varies by tissue, strain, sex, and feeding status (Baar et al. 2016). Microglia from aged mice upregulated mTOR-downstream signaling, affecting targets that regulate translation, including 4EBP1 and pS6 (Keane et al. 2021). We show that phospho-S6 is significantly reduced (~25%) in microglia in response to Cana treatment in the hypothalamus and hippocampus of male mice. However, in female mice reduced phospho-S6, although significant, was detected only in the hypothalamus and reached ~8%. Cana-mediated increases in lifespan may be associated with decreased mTOR signaling.

Cana treatment was associated with improvements in locomotor activity (rotarod), and overall behavior function (open field test). These effects were seen in Cana-treated males, but not in female mice. Previous studies have reported deterioration in a battery of behavioral tests in C57BL/6 mice from 2 to 12 months of age (Shoji *et al*. 2016). In the current study, we show that Cana treatment of genetically diverse UM-HET3 mice significantly improved rotarod performance in male mice, but did not affect grip strength. Additionally, we found that the distance traveled for Cana-treated male mice in the open field test significantly increased, suggesting improved overall activity levels at 30 months of age. Accordingly, Cana-treated aged male mice spent significantly less time at the wall area and more at the center of the maze suggesting the willingness to explore a new environment (Seibenhener & Wooten 2015; Shoji *et al*. 2016; Perals Bertomeu *et al*. 2017). Our results imply that Cana can be beneficial for delaying frailty and may affect more than one system.

In summary, we show that Cana, with recently identified anti-aging properties, can alleviate age-related neuroinflammation, improve central insulin sensitivity and increase overall physical activity. Our findings are in line with other recent clinical studies and rodent work, indicating that Cana and other SGLT2i can improve kidney, heart function, and delay oncogenesis. Our data suggest possible new applications for Cana that require wide-ranging evaluation of patients already receiving this FDA-approved drug for specific indications.

## Materials and Methods

### Animals

These experiments used mice of the UM-HET3 genetically heterogeneous stock as before (Miller *et al*. 2020). Mice were assigned to Control or treatment (Cana) group using a random number table. All the animals were fed Purina 5LG6 until 7 months of age. Mice in the Cana group received this agent at 180 mg per kg of chow, from 7 months of age; this dose is equivalent to 30 milligrams per kg mouse body weight for a 30-gram mouse eating 5 grams of food each day (Miller *et al*. 2020). Animals were maintained under temperature and light-controlled conditions. Procedures involved in this study were approved by the University of Michigan Committee on the Use and Care of Animals (UCUCA).

### Insulin stimulation

On the day of insulin treatment, animals were briefly fasted during the light cycle for <2 hr (to avoid metabolic stress in old mice (Carper *et al*. 2020)), followed by administration of insulin (3 U/kg) i.p. or an equivalent volume of saline. 15 min post injection, animals were anesthetized and perfused as detailed below.

### Perfusion and immunolabeling

Mice were anesthetized and perfused using phosphate buffer saline (PBS) (pH 7.5) followed by 4% paraformaldehyde (PFA). Brains were post-fixed, dehydrated, and sectioned coronally (30μm) using a sliding microtome, followed by immunofluorescent analysis as described (Sadagurski *et al*. 2017). For immunohistochemistry brain sections were washed with PBS six times, blocked with 0.3% Triton X-100 and 3% normal donkey serum in PBS for 2h; then the staining was carried out with the following primary antibodies overnight: rabbit anti-GFAP (1:1000; Millipore, #ab5804), rabbit anti FoxO1 (1:200; Cell signaling #ab2880s) and mouse anti-NeuN (1:1000 Cell Signaling #MAB377). For goat anti-Iba1 (1:1000 Abcam #ab5076) and rabbit anti-TMEM (1:300; Abcam. #ab209064), rabbit anti pS6 (1:100; Cell signaling #2215) immunostaining brain sections were pretreated with 0.5% NaOH and 0.5% H_2_O_2_ in PBS for 20 min. After the primary antibody, brain sections were incubated with AlexaFluor-conjugated secondary antibodies for 2h (Invitrogen). Microscopic images of the stained sections were obtained using an Olympus FluoView 500 and Laser Scanning Confocal Microscope Zeiss LSM 800.

### Quantification

For the quantification of immunoreactivity, images of matched brain areas were taken from at least 3 sections containing the hypothalamus for each brain between bregma −0.82 mm to −2.4 mm (according to the Franklin mouse brain atlas). Serial brain sections were made at 30 μm thickness, and every five sections were represented by one section with staining and cell counting. All sections were arranged from rostral to caudal to examine the distribution of labeled cells. GFAP, Iba1 positive cells, and FoxO1 and pS6 protein expression were counted using ImageJ with DAPI (nuclear staining) and NeuN (neuronal marker). The average of the total number of cells/field was assessed by statistical analysis as detailed below.

### Cell sorting

Microglia and astrocytes were isolated by density gradient from control and Cana-treated mice. This was followed by cell sorting via staining with TMEM119 and CD45/CD11b to identify microglia (CD11b^+^/CD45^low^), characteristics of resident brain microglia, and distinguishing them from CD45^hi^ systemic macrophage populations. Astrocytes were sorted by ACSA2 (Kantzer *et al*. 2017). To validate the sorting we performed the qPCR with sorted microglia and astrocytes using GFAP (astrocytes specific marker) and TMEM119 (microglia specific marker).

### Quantitative Real-Time PCR

Total RNA was isolated from cell sorted microglia and astrocytes using Trizol reagent (Invitrogen, #15596026). The concentration of 1000 ng of RNA was used for cDNA synthesis using a High Capacity cDNA Reverse Transcription Kit (BioRad, #1708891). To detect the contaminated DNA we used the samples processed without the transcriptase reverse enzyme as negative controls. Quantitative real-time PCR was performed using the Applied Biosystems 7500 Real-Time PCR System (Supplementary Table 1, for primers sequence). Each PCR reaction was performed in duplicate. As negative controls, we used water instead of the cDNA, and ß-actin was measured in each cDNA sample as the housekeeping gene. The ΔΔCT method was used to determine the gene transcripts in each sample. For each sample, the threshold cycle (CT) was measured and normalized to the average of the housekeeping gene (ΔCT = CT gene of interest - CT housekeeping gene). The fold change of mRNA in the rest of the samples relative to the control male group was determined by 2-ΔΔCT, where ΔΔCT = ΔCT other animal groups (Cana males, control females, Cana females) - ΔCT control male group. Data are shown as mRNA expression levels relative to the control males.

### Rotarod and grip strength tests

Animals were tested for their ability to balance on an accelerating rotarod at 22 months of age. Animals were placed on the rotarod and the trial began with the spindle revolving at 5 revolutions per minute (RPM) and increased to 40 RPM gradually over a 5-min period. The time at which the animal fell off the rotarod was used as a score, with each animal tested three times and the mean score used in the analysis. Animals were also tested for grip strength using an EB1-BIO-GT3 grip strength meter with an EB1-GRIP-Mouse Grid. Subjects were removed from their cage by the base of the tail and suspended above the grid until their forepaws griped the grid. The tail was gently pulled in a horizontal direction away from the grid until the mouse released its grip. The maximal force was recorded. Each animal was tested six times with a 10 s rest between each. The mean of the six tests was used for analysis. All tests were conducted by an experimenter blind to the treatment group.

### Open field maze

The open-field arena was 23 cm in radius and made with polyvinyl chloride (PVC). The center of the arena was defined as a 10 cm radius circle from the center and outside of this center circle is defined as the outer area (wall area) of the arena. Testing was conducted under bright light. Mice were tested for 10 min inside the arena. At the beginning of the test, animals were placed in the center of the open field arena and the movement of animals was recorded with a Nikon camera. Arenas were cleaned with 70% isopropyl alcohol between trials and the observer was in a different room to the arena throughout the testing time. After recording, the videos were analyzed using Any-Maze Video Tracking System© v 6.34 to analyze the distance moved, the percent time spent in the center of the open field arena, and the percent time spent in the wall area of the arena in each trial. Using Any-Maze Video Tracking System© v 6.34 a series of 20, 40, 60, 80, 100, 120, 140, 160, 180, 200, 220 mm radius zones were identified and used to evaluate the animal track and collect the data for the locomotor activity. The six center circular zones were identified as inner zone and the other six circular zones were identified as outer zone.

### Statistical analysis

Data sets were analyzed using two-factor analysis of variance (two-way ANOVA), using the general linear model function and a full factorial model, which included an effect of treatment (comparing control to Cana treatment), an effect of sex (male or female), and an interaction effect between sex and treatment followed by Newman-Keuls post hoc test. All data were presented as mean ± SEM. p < 0.05 was considered significant. TIBCO Statistica^®^ v. 13.5.0.17 was used for statistical analysis.

## Author contribution

HSMJ, LKD and JJ carried out the research. RAM provided drug-treated and control mice, and reviewed and revised the manuscript. MS designed the study, analyzed the data, wrote the manuscript, and is responsible for the integrity of this work. All authors approved the final version of the manuscript.

## Funding

This study was supported by American Diabetes Association grant #1-lB-IDF-063, Impetus grant, and Michigan Diabetes Research Center P30-DK020572 for MS. Work in the Miller lab was supported by the Glenn Foundation for Medical Research, as well as AG022303 and AG024824

## Disclosure

No conflicts of interest are declared by the authors.

## Data Availability Statement

The data that support the findings of this study are available from the corresponding author upon reasonable request.

## Abbreviations

Cana: Canagliflozin
ACA: acarbose
17aE2: 17-a-estradiol
CNS: central nervous system
ERα: estrogen receptor alpha
ARC: arcuate nucleus of the hypothalamus

---

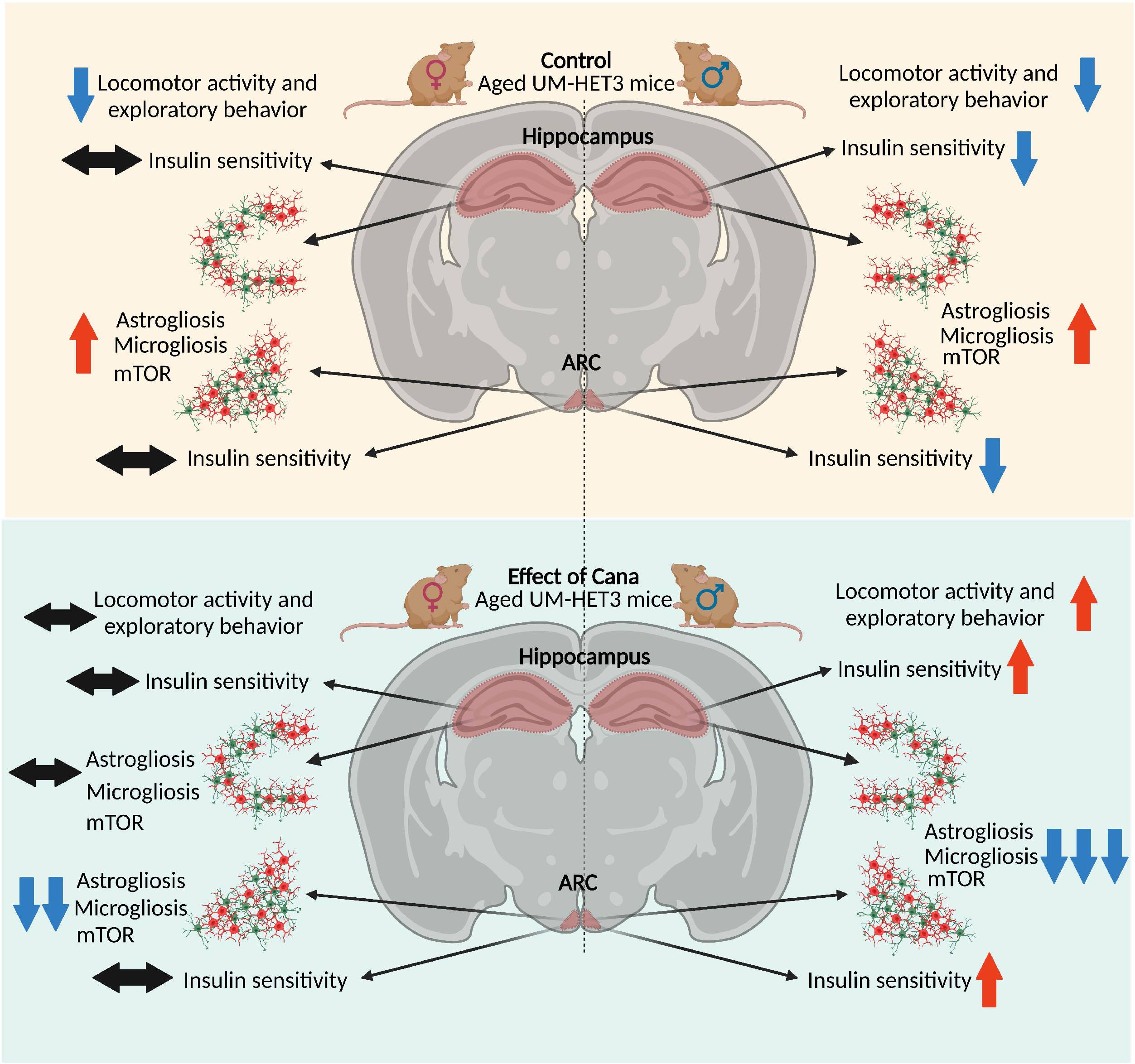

## SUPPLEMENTARY MATERIAL

### Supplementary Legends

**Supplementary Figure 1.**
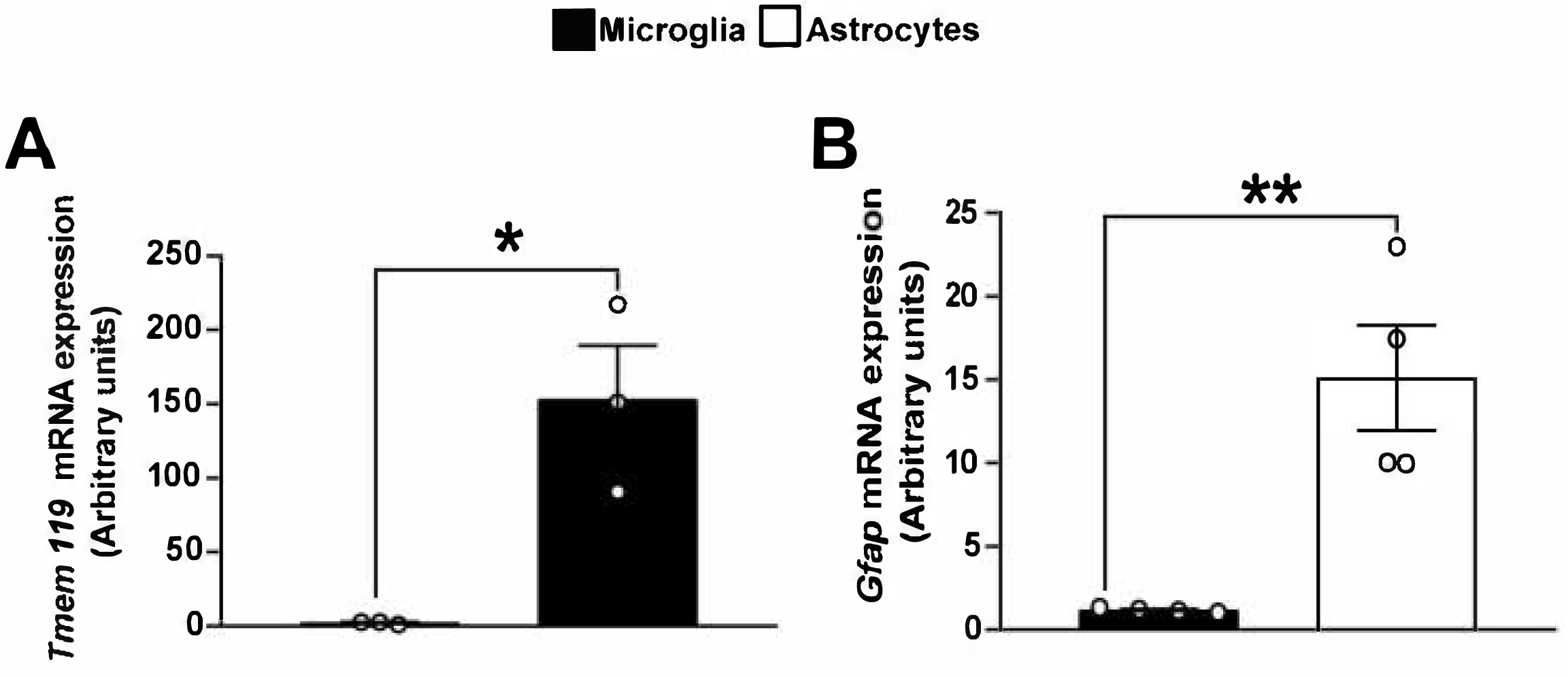
Microglia and astrocytes gene expression. Gene expression of microglia-specific, *Tmem119* (A) and astrocyte-specific, *Gfap* (B). Error bars show SEM for n = 3-4 mice/group. Student’s t-test (*p < 0.05, **p < 0.01).

**Supplementary Figure 2.**
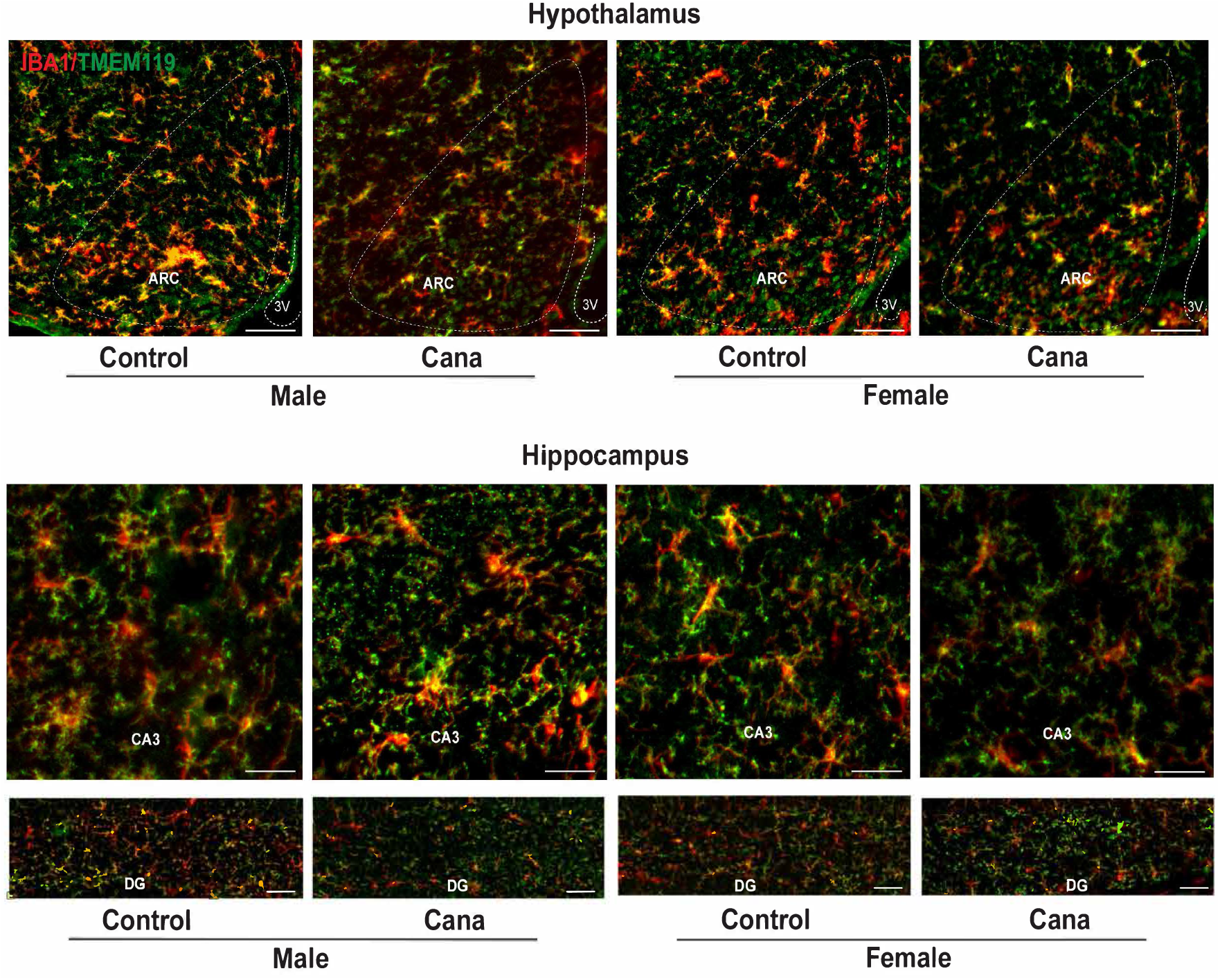
Co-localization of microglial markers Iba1 and TMEM119 in the hypothalamus and hippocampus of Cana-treated mice. Representative images of Iba1 (red) and TMEM119 (green) in the arcuate nucleus of the hypothalamus (ARC) and hippocampal CA3 and dentate gyrus (DG) of 26-28 months-old control and Cana treated mice. Scale bars: 200 μm, 3V, third ventricle.

**Supplementary Figure 3.**
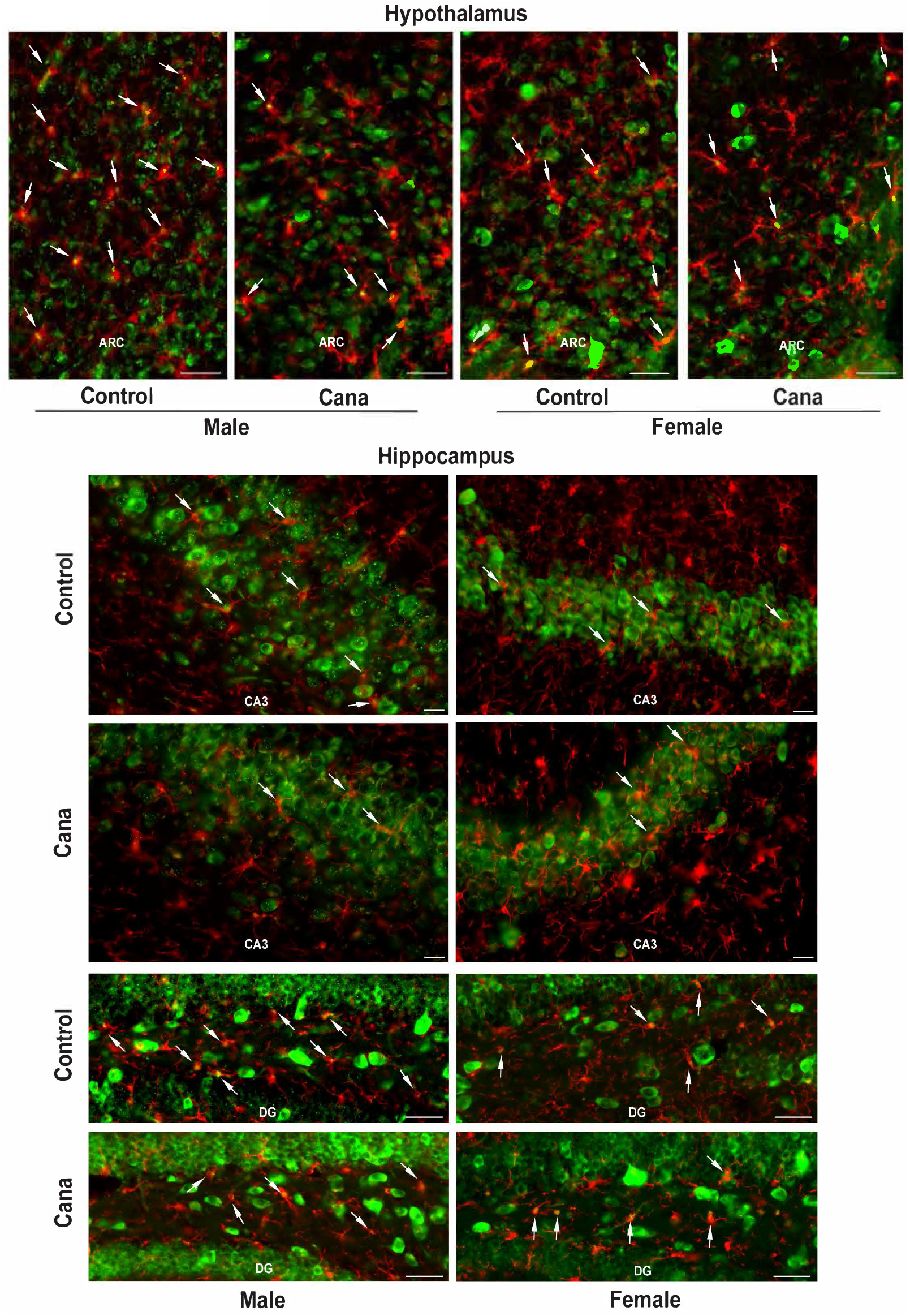
pS6 expression in Cana-treated mice. Representative images of phosphorylated S6 (pS6, green) expression and Iba^+^ cells (red) in the arcuate nucleus of the hypothalamus (ARC) (20x) and hippocampal CA3 and dentate gyrus (DG) (40x) of 26-28-months-old male and female control and Cana-treated mice. Arrows indicate colocalization. Scale bars: 200μm, 3V, third ventricle.

**Supplementary Table 1:**
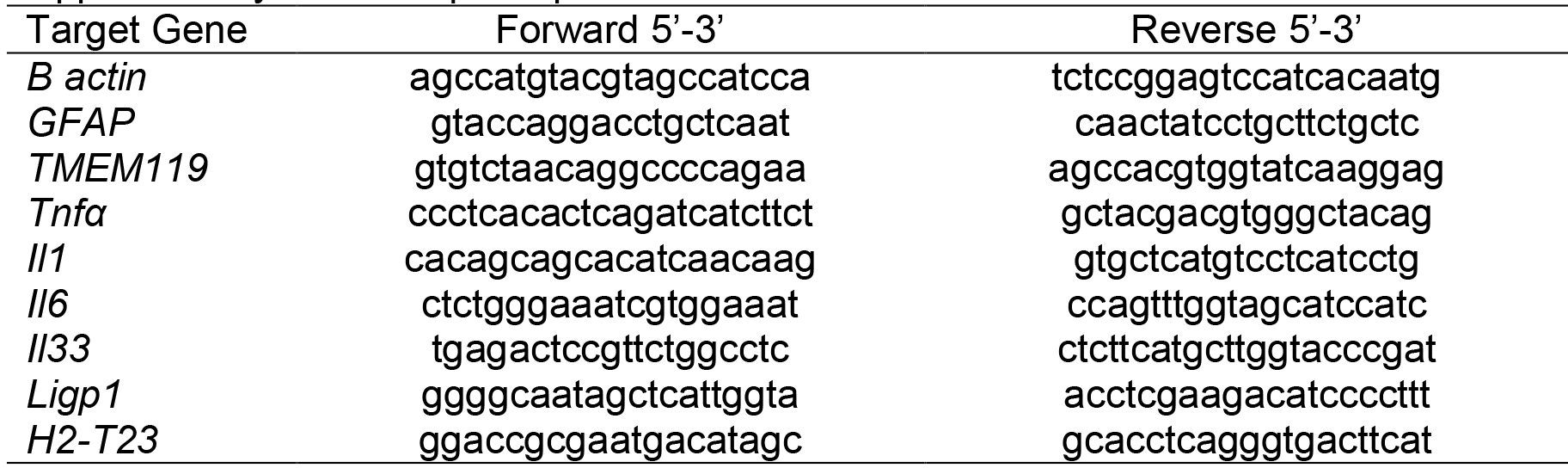
qPCR primer list

## Notes

### Competing Interest Statement

The authors have declared no competing interest.

## REFERENCES

Baar EL, Carbajal KA, Ong IM, Lamming DW (2016). Sex-and tissue-specific changes in mTOR signaling with age in C57BL/6J mice. Aging Cell. 15, 155–166.

Barrientos RM, Kitt MM, Watkins LR, Maier SF (2015). Neuroinflammation in the normal aging hippocampus. Neuroscience. 309, 84–99.

Carlock C, Wu J, Shim J, Moreno-Gonzalez I, Pitcher MR, Hicks J, Suzuki A, Iwata J, Quevado J, Lou Y (2017). Interleukin33 deficiency causes tau abnormality and neurodegeneration with Alzheimer-like symptoms in aged mice. Transl Psychiatry. 7, e1164.

Carper D, Coué M, Laurens C, Langin D, Moro C (2020). Reappraisal of the optimal fasting time for insulin tolerance tests in mice. Molecular Metabolism. 42, 101058.

Clarke LE, Liddelow SA, Chakraborty C, Munch AE, Heiman M, Barres BA (2018). Normal aging induces A1-like astrocyte reactivity. Proc Natl Acad Sci U S A. 115, E1896–E1905.

Clegg DJ, Riedy CA, Smith KAB, Benoit SC, Woods SC (2003). Differential Sensitivity to Central Leptin and Insulin in Male and Female Rats. Diabetes. 52, 682–687.

Fan X, Chan O, Ding Y, Zhu W, Mastaitis J, Sherwin R (2015). Reduction in SGLT1 mRNA Expression in the Ventromedial Hypothalamus Improves the Counterregulatory Responses to Hypoglycemia in Recurrently Hypoglycemic and Diabetic Rats. Diabetes. 64, 3564–3572.

Freude S, Schilbach K, Schubert M (2009). The role of IGF-1 receptor and insulin receptor signaling for the pathogenesis of Alzheimer’s disease: from model organisms to human disease. Curr Alzheimer Res. 6, 213–223.

Furman D, Campisi J, Verdin E, Carrera-Bastos P, Targ S, Franceschi C, Ferrucci L, Gilroy DW, Fasano A, Miller GW, Miller AH, Mantovani A, Weyand CM, Barzilai N, Goronzy JJ, Rando TA, Effros RB, Lucia A, Kleinstreuer N, Slavich GM (2019). Chronic inflammation in the etiology of disease across the life span. Nat Med. 25, 1822–1832.

Garratt M, Bower B, Garcia GG, Miller RA (2017). Sex differences in lifespan extension with acarbose and 17-alpha estradiol: gonadal hormones underlie male-specific improvements in glucose tolerance and mTORC2 signaling. Aging Cell. 16, 1256–1266.

Garratt M, Leander D, Pifer K, Bower B, Herrera JJ, Day SM, Fiehn O, Brooks SV, Miller RA (2019). 17-alpha estradiol ameliorates age-associated sarcopenia and improves late-life physical function in male mice but not in females or castrated males. Aging Cell. 18, e12920.

Głuchowska K, Pliszka M, Szablewski L (2021). Expression of glucose transporters in human neurodegenerative diseases. Biochemical and Biophysical Research Communications. 540, 8–15.

Gonzalez-Freire M, Diaz-Ruiz A, Hauser D, Martinez-Romero J, Ferrucci L, Bernier M, de Cabo R (2020). The road ahead for health and lifespan interventions. Ageing Research Reviews. 59, 101037.

Hallschmid M, Benedict C, Schultes B, Fehm HL, Born J, Kern W (2004). Intranasal insulin reduces body fat in men but not in women. Diabetes. 53, 3024–3029.

Harrison DE, Strong R, Allison DB, Ames BN, Astle CM, Atamna H, Fernandez E, Flurkey K, Javors MA, Nadon NL, Nelson JF, Pletcher S, Simpkins JW, Smith D, Wilkinson JE, Miller RA (2014). Acarbose, 17-alpha-estradiol, and nordihydroguaiaretic acid extend mouse lifespan preferentially in males. Aging Cell. 13, 273–282.

Hierro-Bujalance C, Infante-Garcia C, Del Marco A, Herrera M, Carranza-Naval MJ, Suarez J, Alves-Martinez P, Lubian-Lopez S, Garcia-Alloza M (2020). Empagliflozin reduces vascular damage and cognitive impairment in a mixed murine model of Alzheimer’s disease and type 2 diabetes. Alzheimers Res Ther. 12, 40.

Jais A, Bruning JC (2017). Hypothalamic inflammation in obesity and metabolic disease. J Clin Invest. 127, 24–32.

Kantzer CG, Boutin C, Herzig ID, Wittwer C, Reiss S, Tiveron MC, Drewes J, Rockel TD, Ohlig S, Ninkovic J, Cremer H, Pennartz S, Jungblut M, Bosio A (2017). Anti-ACSA-2 defines a novel monoclonal antibody for prospective isolation of living neonatal and adult astrocytes. Glia. 65, 990–1004.

Keane L, Antignano I, Riechers S-P, Zollinger R, Dumas AA, Offermann N, Bernis ME, Russ J, Graelmann F, McCormick PN, Esser J, Tejera D, Nagano A, Wang J, Chelala C, Biederbick Y, Halle A, Salomoni P, Heneka MT, Capasso M (2021). mTOR-dependent translation amplifies microglia priming in aging mice. The Journal of Clinical Investigation. 131.

Kodali M, Attaluri S, Madhu LN, Shuai B, Upadhya R, Gonzalez JJ, Rao X, Shetty AK (2021). Metformin treatment in late middle age improves cognitive function with alleviation of microglial activation and enhancement of autophagy in the hippocampus. Aging Cell. 20, e13277.

Komleva Y, Chernykh A, Lopatina O, Gorina Y, Lokteva I, Salmina A, Gollasch M (2020). Inflamm-Aging and Brain Insulin Resistance: New Insights and Role of Life-style Strategies on Cognitive and Social Determinants in Aging and Neurodegeneration. Front Neurosci. 14, 618395.

Kullmann S, Hummel J, Wagner R, Dannecker C, Vosseler A, Fritsche L, Veit R, Kantartzis K, Machann J, Birkenfeld AL, Stefan N, Haring HU, Peter A, Preissl H, Fritsche A, Heni M (2021). Empagliflozin Improves Insulin Sensitivity of the Hypothalamus in Humans With Prediabetes: A Randomized, Double-Blind, Placebo-Controlled, Phase 2 Trial. Diabetes Care.

Lam KS, Tiu SC, Tsang MW, Ip TP, Tam SC (1998). Acarbose in NIDDM patients with poor control on conventional oral agents. A 24-week placebo-controlled study. Diabetes Care. 21, 1154–1158.

Lamming DW, Ye L, Katajisto P, Goncalves MD, Saitoh M, Stevens DM, Davis JG, Salmon AB, Richardson A, Ahima RS, Guertin DA, Sabatini DM, Baur JA (2012). Rapamycin-induced insulin resistance is mediated by mTORC2 loss and uncoupled from longevity. Science. 335, 1638–1643.

Liddelow S, Guttenplan K, Clarke L, Bennett F, Bohlen C, Schirmer L, Bennett M, Munch A, Chung W-S, Peterson T, Wilton D, Frouin A, Napier B, Panicker N, Kumar M, Buckwalter M, Rowitch D, Dawson V, Dawson T, Barres B (2017). Neurotoxic reactive astrocytes are induced by activated microglia. Nature. 541.

Lin B, Koibuchi N, Hasegawa Y, Sueta D, Toyama K, Uekawa K, Ma M, Nakagawa T, Kusaka H, Kim-Mitsuyama S (2014). Glycemic control with empagliflozin, a novel selective SGLT2 inhibitor, ameliorates cardiovascular injury and cognitive dysfunction in obese and type 2 diabetic mice. Cardiovasc Diabetol. 13, 148.

Miller RA, Harrison DE, Allison DB, Bogue M, Debarba L, Diaz V, Fernandez E, Galecki A, Garvey WT, Jayarathne H, Kumar N, Javors MA, Ladiges WC, Macchiarini F, Nelson J, Reifsnyder P, Rosenthal NA, Sadagurski M, Salmon AB, Smith DL, Jr., Snyder JM, Lombard DB, Strong R (2020). Canagliflozin extends life span in genetically heterogeneous male but not female mice. JCI Insight. 5.

Milstein JL, Ferris HA (2021). The brain as an insulin-sensitive metabolic organ. Molecular Metabolism. 52, 101234.

Nadon NL, Strong R, Miller RA, Nelson J, Javors M, Sharp ZD, Peralba JM, Harrison DE (2008). Design of aging intervention studies: the NIA interventions testing program. Age (Dordr). 30, 187–199.

Naznin F, Sakoda H, Okada T, Tsubouchi H, Waise TMZ, Arakawa K, Nakazato M (2017). Canagliflozin, a sodium glucose cotransporter 2 inhibitor, attenuates obesity-induced inflammation in the nodose ganglion, hypothalamus, and skeletal muscle of mice. European Journal of Pharmacology. 794, 37–44.

Nguyen T, Wen S, Gong M, Yuan X, Xu D, Wang C, Jin J, Zhou L (2020). Dapagliflozin Activates Neurons in the Central Nervous System and Regulates Cardiovascular Activity by Inhibiting SGLT-2 in Mice. Diabetes Metab Syndr Obes. 13, 2781–2799.

Ohgaki R, Wei L, Yamada K, Hara T, Kuriyama C, Okuda S, Ueta K, Shiotani M, Nagamori S, Kanai Y (2016). Interaction of the Sodium/Glucose Cotransporter (SGLT) 2 inhibitor Canagliflozin with SGLT1 and SGLT2. J Pharmacol Exp Ther. 358, 94–102.

Osataphan S, Macchi C, Singhal G, Chimene-Weiss J, Sales V, Kozuka C, Dreyfuss JM, Pan H, Tangcharoenpaisan Y, Morningstar J, Gerszten R, Patti ME (2019). SGLT2 inhibition reprograms systemic metabolism via FGF21-dependent and-independent mechanisms. JCI Insight. 4.

Perals Bertomeu D, Griffin A, Bartomeus I, Sol D (2017). Revisiting the open-field test: what does it really tell us about animal personality? Animal Behaviour. 123.

Poppe R, Karbach U, Gambaryan S, Wiesinger H, Lutzenburg M, Kraemer M, Witte OW, Koepsell H (1997). Expression of the Na+-D-glucose cotransporter SGLT1 in neurons. J Neurochem. 69, 84–94.

Sadagurski M, Cady G, Miller RA (2017). Anti-aging drugs reduce hypothalamic inflammation in a sexspecific manner. Aging Cell. 16, 652–660.

Seibenhener ML, Wooten MC (2015). Use of the Open Field Maze to measure locomotor and anxiety-like behavior in mice. Journal of visualized experiments: JoVE, e52434–e52434.

Shoji H, Takao K, Hattori S, Miyakawa T (2016). Age-related changes in behavior in C57BL/6J mice from young adulthood to middle age. Mol Brain. 9, 11–11.

Sierra A, Gottfried-Blackmore AC, McEwen BS, Bulloch K (2007). Microglia derived from aging mice exhibit an altered inflammatory profile. Glia. 55, 412–424.

Spallone V, Valensi P (2021). SGLT2 inhibitors and the autonomic nervous system in diabetes: A promising challenge to better understand multiple target improvement. Diabetes & Metabolism. 47, 101224.

Strong R, Miller RA, Antebi A, Astle CM, Bogue M, Denzel MS, Fernandez E, Flurkey K, Hamilton KL, Lamming DW, Javors MA, de Magalhaes JP, Martinez PA, McCord JM, Miller BF, Muller M, Nelson JF, Ndukum J, Rainger GE, Richardson A, Sabatini DM, Salmon AB, Simpkins JW, Steegenga WT, Nadon NL, Harrison DE (2016). Longer lifespan in male mice treated with a weakly estrogenic agonist, an antioxidant, an alpha-glucosidase inhibitor or a Nrf2-inducer. Aging Cell. 15, 872–884.

Tahara A, Takasu T, Yokono M, Imamura M, Kurosaki E (2016). Characterization and comparison of sodium-glucose cotransporter 2 inhibitors in pharmacokinetics, pharmacodynamics, and pharmacologic effects. J Pharmacol Sci. 130, 159–169.

Talbot K, Wang H-Y, Kazi H, Han L-Y, Bakshi KP, Stucky A, Fuino RL, Kawaguchi KR, Samoyedny AJ, Wilson RS, Arvanitakis Z, Schneider JA, Wolf BA, Bennett DA, Trojanowski JQ, Arnold SE (2012). Demonstrated brain insulin resistance in Alzheimer’s disease patients is associated with IGF-1 resistance, IRS-1 dysregulation, and cognitive decline. The Journal of Clinical Investigation. 122, 1316–1338.

von Bernhardi R, Eugenin-von Bernhardi L, Eugenin J (2015). Microglial cell dysregulation in brain aging and neurodegeneration. Front Aging Neurosci. 7, 124.

Wang Y, Fu WY, Cheung K, Hung KW, Chen C, Geng H, Yung WH, Qu JY, Fu AKY, Ip NY (2021). Astrocyte-secreted IL-33 mediates homeostatic synaptic plasticity in the adult hippocampus. Proc Natl Acad Sci U S A. 118.

Zhurova EA, Zhurov VV, Chopra D, Stash AI, Pinkerton AA (2009). 17Alpha-estradiol x 1/2 H2O: super-structural ordering, electronic properties, chemical bonding, and biological activity in comparison with other estrogens. Journal of the American Chemical Society. 131, 17260–17269.

